# Zika infection of neural progenitor cells perturbs transcription in neurodevelopmental pathways

**DOI:** 10.1101/072439

**Authors:** Lynn Yi, Harold Pimentel, Lior Pachter

## Abstract

**Background:** A recent study of the gene expression patterns of Zika virus (ZIKV) infected human neural progenitor cells (hNPCs) revealed transcriptional dysregulation and identified cell-cycle-related pathways that are affected by infection. However deeper exploration of the information present in the RNA-Seq data can be used to further elucidate the manner in which Zika infection of hNPCs affects the transcriptome, refining pathway predictions and revealing isoform-specific dynamics.

**Methodology/Principal Findings:** We analyzed data published by Tang *et al.* using state-of-the-art tools for transcriptome analysis. By accounting for the experimental design and estimation of technical and inferential variance we were able to pinpoint Zika infection affected pathways that highlight Zika’s neural tropism. The examination of differential genes reveals cases of isoform divergence.

**Conclusions/Significance:** Transcriptome analysis of Zika infected hNPCs has the potential to identify the molecular signatures of Zika infected neural cells. These signatures may be useful for diagnostics and for the resolution of infection pathways that can be used to harvest specific targets for further study.

## Introduction

As infection with Zika virus (ZIKV) is associated with increasing cases of congenital microcephaly and adult Guillain-Barre Syndrome, a characterization of its pathophysiology becomes crucial. A molecular characterization of the effects of infection may help in the development of fetal diagnostics and can accelerate the identification of crucial genes and pathways critical in disease progression. RNA-Sequencing (RNA-Seq) is an effective technology for probing the transcriptome and has been applied to study the effects of ZIKV infection of human neuroprogenitor cells (hNPCs) [1].

While initial analyses of the data have been used to conduct a general survey of transcriptome changes upon infection [1–3], they [1,2] used a method, Cufflinks/Cuffdiff [4], that fail to take advantage of the experimental design used in Tang et. al [1]. They [1,2,3] also do not examine transcriptome dynamics at the isoform level.

We apply the recently developed kallisto [5] and sleuth [6] programs to improve the accuracy of quantification and to extract information from the data that was previously inaccessible. We find that sleuth’s improved control of false discovery rate results in differential transcript and gene lists that are much more specific and that are significantly enriched in neurodevelopmental pathways. They reveal ZIKV’s neural tropism and the host’s response to viral infection. Furthermore, we demonstrate that the combination of accurate kallisto quantification, assessment of inferential variance and the sleuth response error model allows for the detection of post infection isoform-specific changes that were missed in previous analyses.

The sleuth Shiny app drives a freely available website that allows for reproducibility of our analyses, and provides tools for interacting with the data. This makes the dataset useful for analysis by infectious disease experts who may not have bioinformatics expertise.

## Methods

We ran kallisto and sleuth on a total of eight samples of ZIKV infected and mock infected hNPCs (GEO: Series GSE78711). The runs were performed on a laptop and can be repeated using the provided scripts at http://www.github.com/pachterlab/zika/. We used kallisto to pseudo-align the RNA-seq reads, building an index using the ENSEMBL GRC38 release 85 *Homo sapiens* transcriptome and using default parameters (kmer size = 31, fragment length = 187 and sd = 70 for the single end reads), quantifying transcript abundances, and performing 100 bootstraps per sample. To identify differentially transcripts and genes we first modified sleuth to be able to take advantage of the technical replicates performed by Tang et. al [1]. This was done by replacing an estimate of inferential variance from an average of bootstrap estimated variances to a weighting based on the number of fragments in each replicate. The response error model of sleuth was then used to identify statistically significant differential genes and transcripts.

**Table 1a:** Experimental design and inferential variance estimation weights

| Sample | Accession Number | Condition | Seq method | Seq machine | Reads | Fragments / weights |
| --- | --- | --- | --- | --- | --- | --- |
| Mock1-1 | SRR3191542 | mock | paired-end | MiSeq | 15855554 | 7927777 |
| Mock2-1 | SRR3191543 | mock | paired-end | MiSeq | 14782152 | 7391076 |
| ZIKV1-1 | SRR3191544 | zika | paired-end | MiSeq | 14723054 | 7361527 |
| ZIKV2-1 | SRR3191545 | zika | paired-end | MiSeq | 15242694 | 7621347 |
| Mock1-2 | SRR3194428 | mock | single-end | NextSeq | 72983243 | 72983243 |
| Mock2-2 | SRR3194429 | mock | single-end | NextSeq | 94729809 | 94729809 |
| ZIKV1-2 | SRR3194430 | zika | single-end | NextSeq | 71055823 | 71055823 |
| ZIKV-2-2 | SRR3194431 | zika | single-end | NextSeq | 66528035 | 66528035 |

## Results

We detected 4610 transcripts across 3646 genes that are differentially expressed between ZIKV and mock infected samples (false discovery rate of 0.05) (Fig 1: principle component analysis. S1 Table: differentially expressed transcripts, sorted by significance level). 2895 of the 3646 differentially expressed genes were also reported in Tang et. al [1], but they report an additional 3969 genes that we failed to find containing a significant transcript (they found a total of 6864 significant genes), whose 18423 transcripts have an average qval = 0.55. Furthermore, we found 751 differentially expressed genes corresponding to 5426 transcripts not detected by Cufflinks.

**Figure 1:**
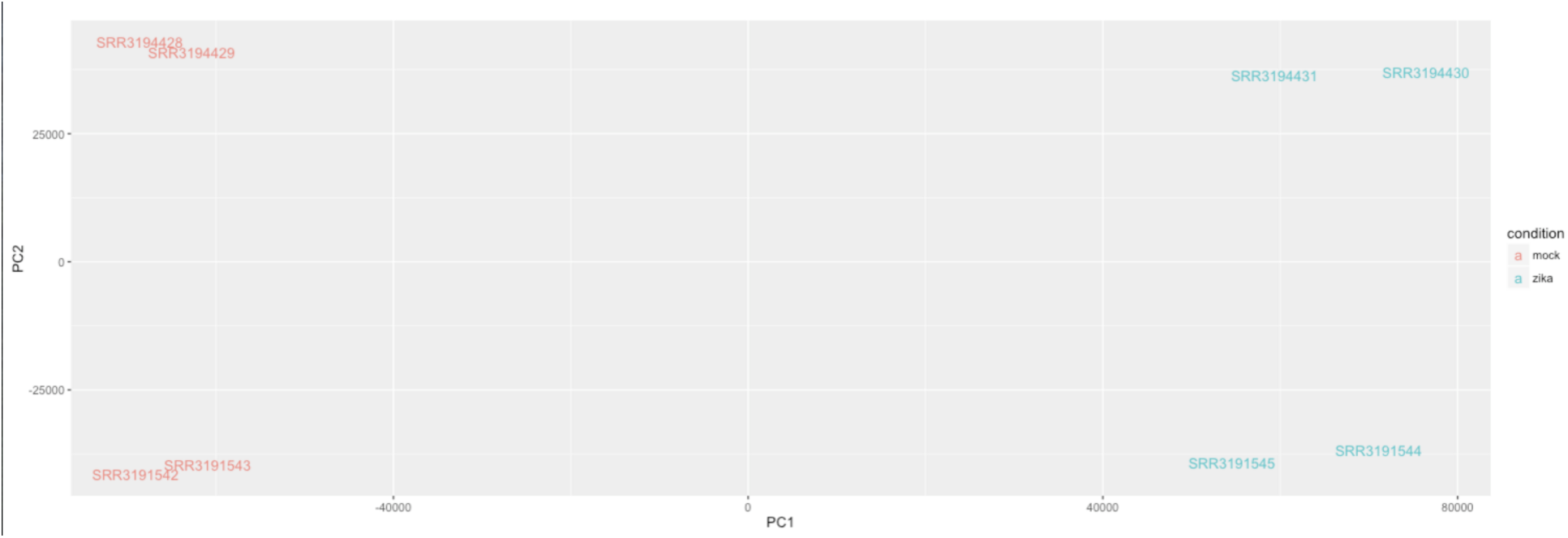
Principle component analysis shows that the primary contributor to variance is whether the sample is ZIKV-infected or mock-infected. The secondary component is method of sequencing, i.e. paired-ends or single-end.

The statistics and figure above, along with interactive data visualization tools, can be found via Sleuth’s Shiny app: http://lair.berkeley.edu/tang16/.

### Zika induced isoform divergence

Differentially regulated genes may be missed in gene-level differential analysis for several reasons. Noise in the measurement of a highly expressed transcript can mask expression changes in lowly expressed transcripts. In the case of isoform switching, the upregulation in one isoform can “cancel out” the effects of downregulation in another. We identified 108 genes that undergo isoform divergence as a result of infection, where isoform divergence is defined as a gene containing one or more transcripts that are are significantly upregulated and at least one other transcript that is significantly downregulated (see S2 Table of isoform diverging transcripts with statistics). Of these 108, 57 genes were not considered differentially expressed genes by Cuffdiff analysis, corresponding to 150 transcripts.

An analysis on these 108 isoform diverging genes using Reactome pathway analysis [7] show pathway enrichment in neuronal system (specifically transmission across chemical synapses and protein-protein interactions at the synapses), developmental biology (specifically axon guidance), immune system, DNA repair, chromatin modifying enzymes, gene expression (rRNA and transcriptional regulation), metabolism, signal transduction, transmembrane transport and vesicle-mediated transport.

One of these 57 isoform diverging genes not picked up by Cufflink is NRCAM, neuronal cell adhesion molecule, which according to Gene Cards is putatively involved in neuron-neuron adhesion and axonal cone growth. Another is CHRNA7, cholinergic receptor nicotinic alpha 7 subunit. [8] See Fig 2a and 2b for plots of the changes in their transcripts levels.

**Figure 2a and b:**
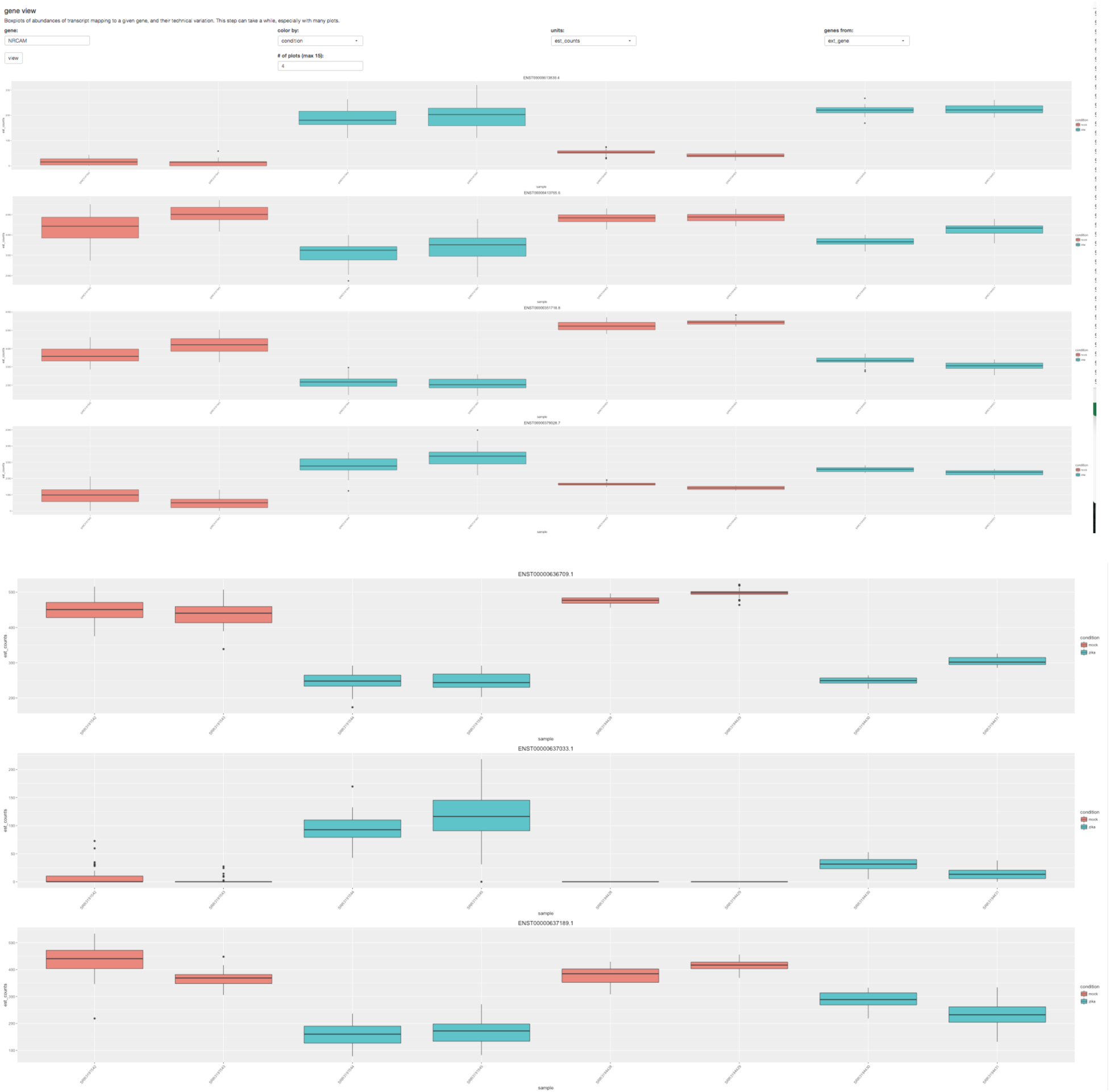
Examples of genes with divergent isoforms, NRCAM and CHRNA7, viewed with our Shiny app. For a specific gene, it displays each transcript and the abundances corresponding to each sample.

### A gene ontology (GO) analysis of sleuth-discovered genes showcase neural and head development networks

We analyzed the set of 3656 genes with differentially regulated transcripts with a gene ontology tool, ClueGO a plugin for Cytoscape [9, 10], over the Biological Processes ontology network, using GO Term Fusion. We set the network specificity to global (GO tree interval: 1-4), using pathways with a minimum of 50 genes and kappa score of 0.5. The enriched nodes of particular interest include neuron projection guidance (pval = 2.7E-3 vs >0.05 with Cuffdiff), cerebral cortex development (1.6E-7 vs >0.05), neuron development (9.9E-6 vs 3.9E-4), neuron projection development (1.8E-6 vs 5.0E-5), nervous system development (3.0E-10 vs 1.0E-9), central nervous system development (6.9E-9 vs 1.0E-4), brain development (2.8E-9 vs 8.0E-4), forebrain development (1.9E-7 vs 4.1E-2), telecephalon development (2.7E-5 vs 5.2E-3), head development (pval = 1.3E-6 vs 3.2E-4), and cellular response to stress (9.4E-26 vs 7.3E-22).

**Figure 3:**
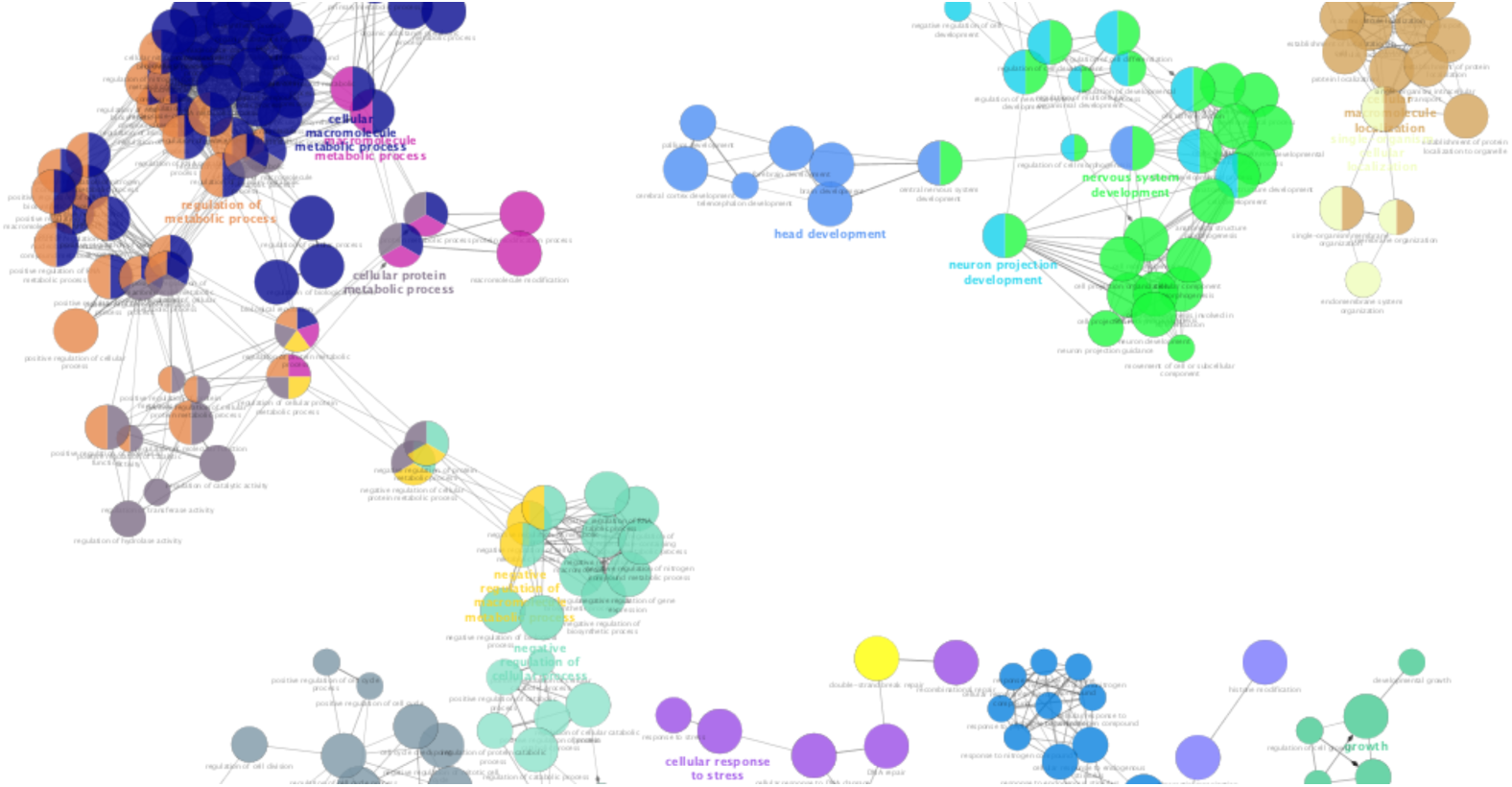
A subgraph of the network resulting from ClueGO analysis of the differentially regulated genes discovered by kallisto and sleuth.

A GO analysis results tables and ctyoscape JSON file (see Supplementary materials) that can be used to render the network in cytoscape.

## Discussion

RNA-Seq can provide rapid and highly resolved probing of infection response, and initial studies of Zika infection dynamics highlight neurally active isoforms, genes and pathways that may play an important role in disease etiology. However the simplicity of RNA-Seq library prep and cDNA sequencing belies the complexity of analysis. We have shown that a careful analysis of previously published data can reveal novel targets with higher confidence, and in the process rendering a valuable dataset usable by the community of Zika researchers.

The kallisto and sleuth tools we have used in our analysis are particularly powerful when coupled with the interactive sleuth Shiny application, and our publicly available server providing access to our analysis contains numerous interactive plots and analyses that cannot be reproduced in a static publication. This highlights the utility and importance of data sharing [11], and we hope that our analysis, aside from its usefulness for the Zika scientific community, can also serve as a blueprint for future data sharing efforts.

sleuth is a fast and accurate pipeline for analyzing RNA-Seq data that allows for testing at the isoform level. The alignment and quantification pipeline is feasible and compatible with a standard desktop computer. The interactive Sleuth application, made publically available, allows for informative data visualization, including those of library prep fragment size distributions, principle component analysis, and gene and transcript expression changes. We invite the scientific community studying Zika to utilize this toolkit.

## References

[1] Tang H, Hammack C, Ogden SC, Wen Z, Qian X, Li Y, Yao B, Shin J, Zhang F, Lee EM, Christian KM, Didier RA, Jin P, Song H, Ming GL, Zika Virus Infects Human Cortial Neuro Progenitors and Attenuate Their Growth. Cell Stem Cell. 2016 May 5;18(5):587–90.

[2] Rolfe AJ, Bosco DB, Wang J, Nowakowski RS, Fan J, Ren, Bioinformatic analysis reveals the expression of unique transcriptomic signatures in Zika virus infected human neural stem cells, Cell Biosci. 2016 6:42, doi: 10.1186/s13578-016-0110-x.

[3] Wang Z, Ma’ayan A, An open RNA-seq data analysis pipeline tutorial with an example of reprocessing data from a recent Zika virus study. F1000Research 2016, 5:1574, doi:10.5256/f1000research.9804.r14924.

[4] C. Trapnell, A. Roberts, L. Goff, G. Pertea, D. Kim, D.R. Kelley, H. Pimentel, S.L. Salzberg, J.L. Rinn and L. Pachter, Differential gene and transcript expression analysis of RNA-seq experiments with TopHat and Cufflinks, Nature Protocols, 7 (2012), 562–578.

[5] Bray NL, Pimentel H, Melsted P and Pachter L, Near-optimal probabilistic RNA-seq quantification, Nature Biotechnology 34, 525–527 (2016), doi:10.1038/nbt.3519.

[6] Pimentel HJ, Bray N, Puente S, Melsted P, Pachter L, Differential analysis of RNA-Seq incorporating quantification uncertainty, bioRxiv 058164; doi: http://dx.doi.org/10.1101/058164.

[7] Croft D, Mundo AF, Haw R, Milacic M, Weiser J, Wu G, Caudy M, Garapati P, Gillespie M, Kamdar MR, Jassal B, Jupe S, Matthews L, May B, Palatnik S,Rothfels K, Shamovsky V, Song H, Williams M, Birney E, Hermjakob H, Stein L, D’Eustachio P. The Reactome pathway knowledgebase. Nucleic Acids Res. 2014 Jan;42(Database issue):D472–7. doi: 10.1093/nar/gkt1102. Epub 2013 Nov 15.

[8] Stelzer G, Rosen R, Plaschkes I, Lieder I, Zimmerman S, Twik M, Fishilevich S, Nudel R, Kohn A, Mazor Y, Kaplan S, Iny Stein T, Warshawsky D, Guan-Golan Y, Rappaport N, Safran M, and Lancet D. The GeneCards Suite: From Gene Data Mining to Disease Genome Sequence Analysis, Current Protocols in Bioinformatics (2016), in press

[9] Bindea G, Mlecnik B, Hackl H, Charoentong P, Tosolini M, et al. ClueGO: a Cytoscape plug-in to decipher functionally grouped gene ontology and pathway annotation networks. Bioinformatics. 2009, 25(8):1091–3.

[10] Bindea G, Galon J, Mlecnik B, CluePedia Cytoscape plugin: pathway insights using integrated experimental and in silico data. Bioinformatics. 2013, 29(5):661–3.

[11] Longo D, Drazen J, Data Sharing. N Engl J Med 2016; 374:276–277, January 21, 2016, DOI: 10.1056/NEJMe1516564.

